# Engineering cell and nuclear morphology on nano topography by contact-free protein micropatterning

**DOI:** 10.1101/2023.06.05.543791

**Authors:** Einollah Sarikhani, Dhivya Pushpa Meganathan, Keivan Rahmani, Ching-Ting Tsai, Abel Marquez-Serrano, Xiao Li, Francesca Santoro, Bianxiao Cui, Lasse Hyldgaard Klausen, Zeinab Jahed

## Abstract

Platforms with nanoscale topography have recently become powerful tools in cellular biophysics and bioengineering. Recent studies have shown that nanotopography affects various cellular processes like adhesion and endocytosis, as well as physical properties such as cell shape.

To engineer nanopillars more effectively for biomedical applications, it is crucial to gain better control and understanding of how nanopillars affect cell and nuclear physical properties, such as shape and spreading area, and impact cellular processes like endocytosis and adhesion. In this study, we utilized a laser-assisted micropatterning technique to manipulate the 2D architectures of cells on 3D nanopillar platforms. We performed a comprehensive analysis of cellular and nuclear morphology and deformation on both nanopillar and flat substrates. Our findings demonstrate precise engineering of cellular architectures through 2D micropatterning on nanopillar platforms. We show that the coupling between nuclear and cell shape is disrupted on nanopillar surfaces compared to flat surfaces. Furthermore, we discovered that cell elongation on nanopillars enhances nanopillar-induced endocytosis. These results have significant implications for various biomedical applications of nanopillars, including drug delivery, drug screening, intracellular electrophysiology, and biosensing. We believe our platform serves as a versatile tool for further explorations, facilitating investigations into the interplay between cell physical properties and alterations in cellular processes.

**Graphical Abstract:** 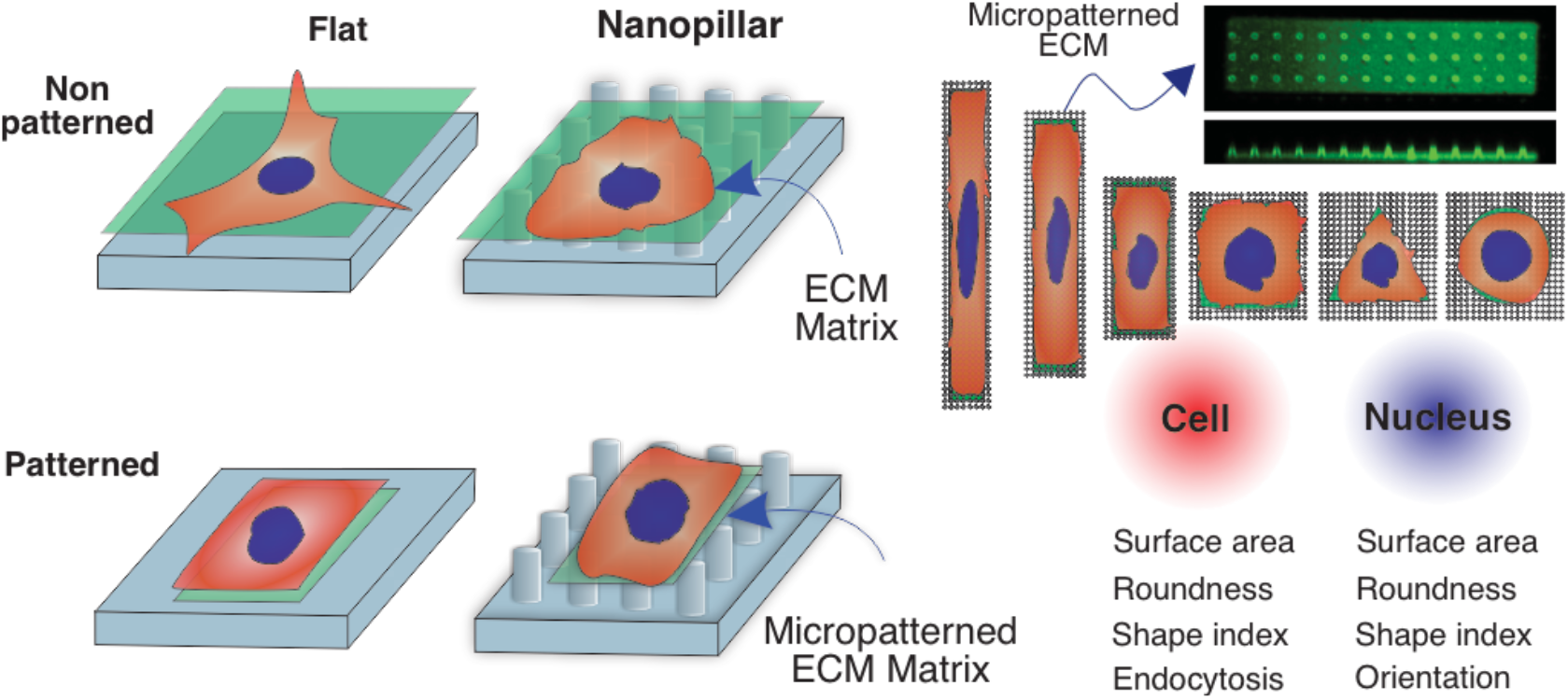

## INTRODUCTION

### Nanopillars are powerful platforms for cellular biophysics and bioengineering

In recent years, platforms with nanoscale topography have become increasingly attractive for supporting cell growth for a wide range of application in areas such as tissue engineering^1–3^, drug delivery, discovery, and development ^4–6^, biosensing^7–9^, electrophysiology^10^, as well as fundamental cell biology, biophysics and mechanobiology. One type of nanostructured platform that has gained significant attention are repeating patterns of free-standing cylinders with diameters of few hundred nanometers commonly known as nanopillar arrays.^11^ Because of the unique properties of nanopillars such as high surface area-to-volume ratio, biocompatibility, controllability of their geometry and material, and the availability of several techniques for their fabrication, nanopillar platforms are being developed for a variety of fundamental research and commercial clinical applications^12^ in the biomedical field.^13^ For example, in recent studies, nanopillars arrays have been used as platforms for biomolecule delivery^14,15^, building blocks of state-of-the-art nanobioelectronics for neuron^16,17^ and cardiac electrophysiology^18–21^, platforms for cancer malignancy detection^22^, and modulation of cell and nuclear mechanics. ^23–25^ Due to the growing list of applications of nanopillars in the biomedical field, several studies have focused on characterizing the interface between nanopillars and cells with the goal of controlling cell behaviors.^26,27^

### Nanopillars alter cellular properties and processes

Earlier studies showed that cells growing on nanopillars exhibit altered behavior compared to the flat surfaces showing changes in several cellular processes including endocytosis^28–30^, adhesion^31^, proliferation^32,33^, migration^32,33^, and differentiation^34–36^. These recent discoveries of nanopillar-induced changes in cell processes have enabled the emergence of the new and exciting applications mentioned above. For example, nanopillars enhance biomolecule delivery efficiency due to their enhancement of endocytosis events.^28,29^ However, more recent studies have shown that nanopillars not only alter cellular processes, but they also change several cell and nuclear biophysical properties such as stiffness, volume, and spreading area. Therefore, achieving a more rational approach to engineering nanopillars for various biomedical applications necessitates a better control and understanding of the interplay between nanopillar-induced changes in cell and nuclear properties such as shape and spreading area, and alterations in cellular processes such as endocytosis and adhesion. To investigate this interplay, we first present a contact-free micropatterning technique for engineering cellular morphologies on nanopillar arrays. We show that properties such as nuclear and cell shape and spreading area can be precisely controlled on nanopillars. We then compared the controlled cellular and nuclear morphologies on flat and nanopillar substrates. We further investigated the effects of substrate topography and cell shape on endocytosis and found that high elongation of cells on nanopillars and on flat substrates can further promote endocytosis. Overall, our results demonstrate that our platform can be used to precisely control cellular and nuclear properties on nanopillars which in turn impact cellular processes. These results, and future studies using the presented technique can lead to a more rational design of nanopillar platforms for various biomedical applications.^46,47^

## RESULTS AND DISCUSSION

### ECM-Micropatterns on transparent SiO_2_ nanopillar arrays

We fabricated fused silica *SiO*_*2*_ nanopillar arrays using photolithography followed by consecutive dry etching and wet etching process **(Figure 1A**) as detailed in the experimental section. The size and shape of the fabricated nanopillars were characterized with scanning electron microscopy as shown in **Figure 1B** and the dimensions were determined (Average height of 1.51 ± 0.19, diameter of 0.41 ± 0.09 μm and pitch of 2.55 ± 0.11). To pattern Extracellular Matrix (ECM) proteins on nanopillar arrays, we used the light-induced molecular adsorption protein (LIMAP) technique with a maskless photolithography system.^37^ This technique is a maskless, controllable and non-contact patterning of protein that accommodates patterning on a variety of substrates such as glass, PDMS and is compatible with our transparent nanopillar arrays.^38,39^ To achieve specific attachment of cells to micropatterns, the surface was first coated with a poly-L-lysine layer through electrostatic interaction and next covalently coated with an “antifouling” reagent, polyethylene glycol (PEG). Next, we deposited a photoinitiatior (PLPP, Alvéole Lab) that cleaves the PEG molecules with high accuracy upon UV light (375 nm) activation. The nanopillar chip was mounted on a cover glass #1 (Fisher brand™) and placed on the Nikon Ti-E microscope stage, followed by UV exposure through a digital micromirror device (DMD) controlled by Leonardo plugin (Alvéole Lab) software with micromanager software v.1.4.22 used to apply various doses of UV.

**Figure 1.**
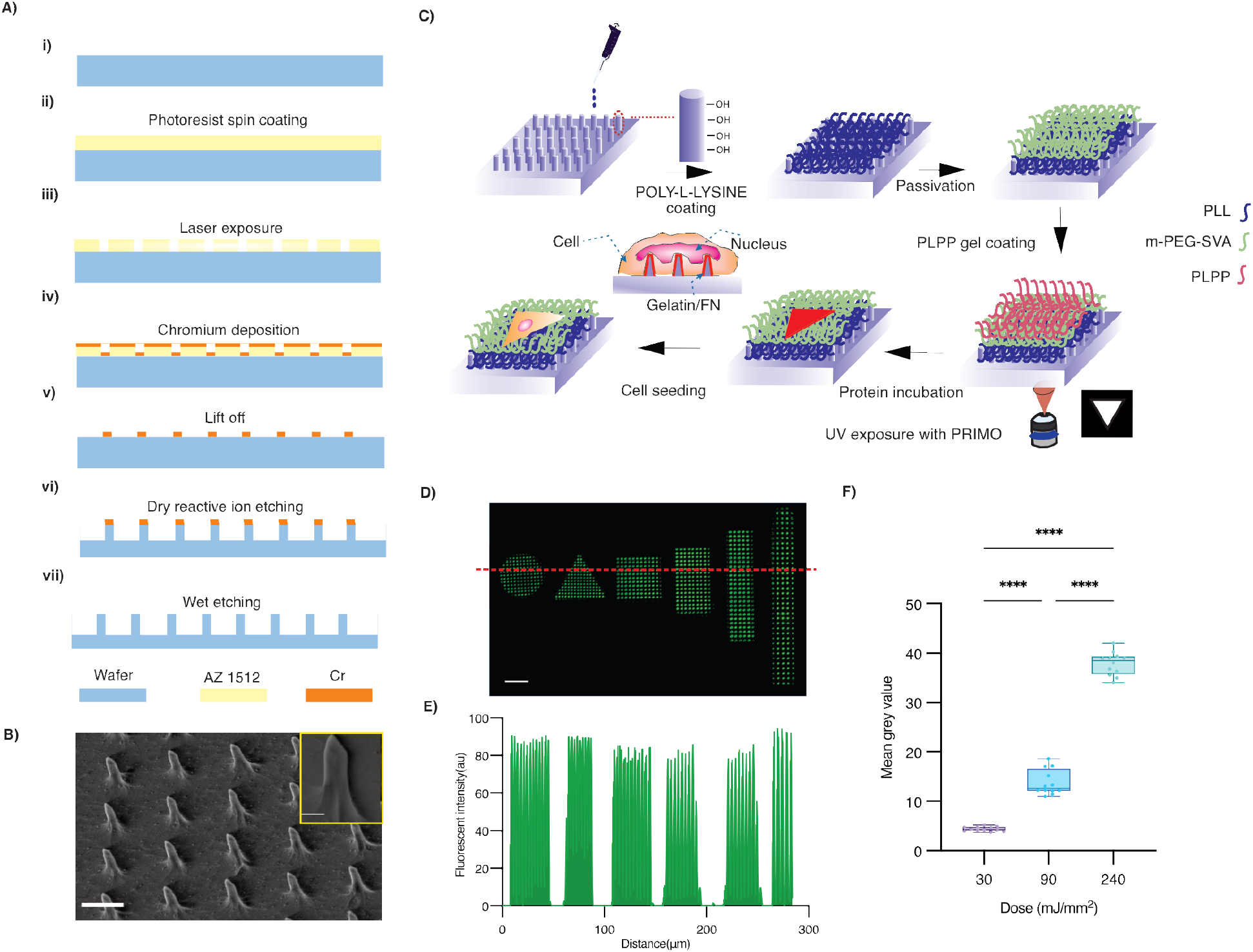
Fabrication of nanopillar substrates and contact-free micropatterning of proteins on nanopillars. A) overview schematic of the nanopillars fabrication process including photolithography, electron beam evaporation, dry etching and wet etching. B) SEM image (45°) of nanopillar arrays with 1.51 ± 0.19 μm height, 0.41 ± 0.09 μm diameter and 2.55 ± 0.11 pitch (scale bar =5 μm). Inset shows an enlarged image of single pillar representing the size and shape of pillars (scale bar =1 μm) C) Schematic of protein photopatterning on the nanopillars with PRIMO system D) FITC-gelatin protein patterned absorbed on the patterned area with various shapes on nanopillars. E) Profile of protein intensity across the red line on image 1D. F) Average fluorescent intensity of FITC-gelatin micropatterns as a function of the UV exposure dose.

Finally, we coated the exposed areas with a mixture of the FITC-gelatin and FN to enhance cellular attachment and visualize the micropatterned areas on the flat surface (**Figure S1**) and nanopillar arrays **(Figure 1C, 1D)**. To obtain the highest contrast and protein adsorption on the micropatterned areas, we optimized the dose of UV illumination by testing UV doses of 30, 90, and 240 mJ/mm2 and obtained the optimized results at 240 mJ/mm2 which demonstrated the highest fluorescent intensity compared to the others (**Figure 1F**) (higher dose increased the time for patterning and patterns resolution decreased). In addition, the fluorescent intensity profile of the protein across a line showed highest intensity on micropatterned areas, and relatively low background intensity between micropatterns. This is essential for promoting specific cell attachment only on micropatterned areas (**Figure 1E**). Using this technique, we robustly and reproducibly patterned hundreds of ECM shapes on nanopillars, and successfully controlled cell morphology on nanopillars as described in the following section.

### Different cell types conform to micropatterns of various shapes on nanopillars

Having established a robust method for developing gelatin/FN micropatterns on nanopillars, we set out to control cellular morphologies on nanopillars by confining single cells to each micropattern. To this end, we tested a variety of cell densities and found an optimal cell seeding density of 15,000 cells/cm^2^ to obtain the highest number of single cells on micropatterns. We cultured cells on micropatterns with a range of shape parameters (roundness, and shape index) (**Figure 2A**) and showed that cells conform to micropatterns of various shapes parameters on nanopillar by fluorescent imaging of cell boundaries by using an actin stain (**Figure 2A**). To demonstrate the versatility of the current method for a variety of cell types, we seeded two cell types (Human Bone Osteosarcoma Epithelial Cells (U2OS) and Human Mesenchymal Stem Cells (hMSC)) and showed qualitatively that both conform to the shapes of micropatterns (**Figure 2A, 2B**).

**Figure 2.**
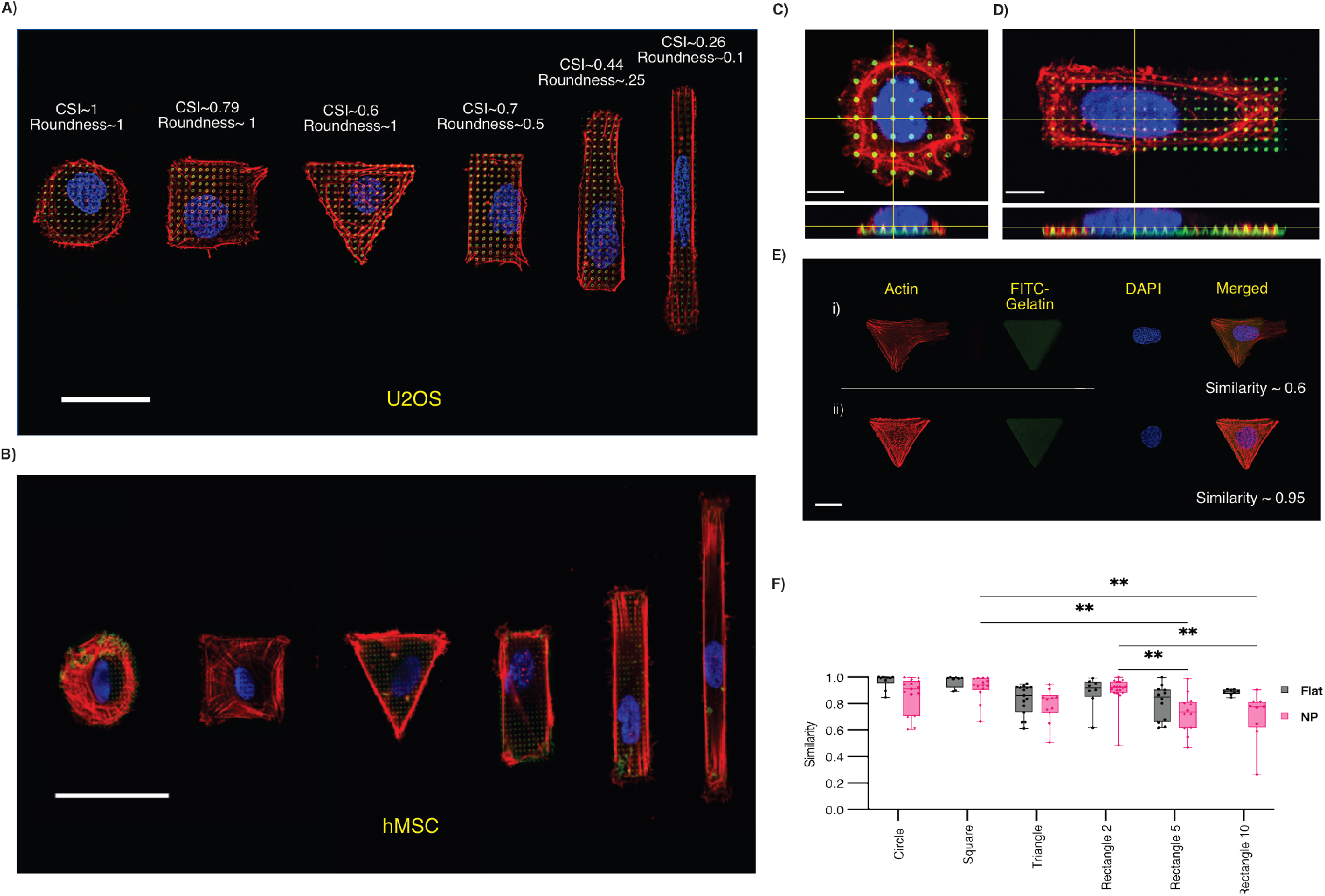
Controlling cell morphology on nanopillars with contact-free micropatterning. A) representative widefield image of U2OS cells on patterned proteins on nanopillars showing conformation of cells to the ECM matrix. (Scale bar = 40 μm) B) representative widefield image of hMSC cells on patterned proteins on nanopillars. ECM protein gelatin (green), Actin (red) and nucleus (blue) are shown. (Scale bar= 50 μm) C) Representative confocal image of U2OS cell on circular micropatterned ECM protein on nanopillars from top and side view. (Scale bar= 10 μm) D) Representative confocal image of U2OS cell on rectangular micropatterned ECM protein on nanopillars from top and side view. (Scale bar= 10 μm) E) Representative images of U2OS cells conforming to the ECM protein with different similarity index. ECM protein gelatin (green), Actin (red) and nucleus (blue) are shown, F) Similarity analysis of the cells conforming to the shape of the ECM proteins patterned on flat and nanopillar substrate. (Scale bar= 20 μm). (**p < 0.01)

To determine whether cells spread in 2D on top of the nanopillar, or wrap around nanopillars in 3D, we performed confocal fluorescence microscopy. Reconstructed side views of the cell-nanopillar interface demonstrate the cells conforming to the micropatterned area while the cell and its nucleus deforms over the nanopillar arrays to engulf the nanopillars (**Figure 2C and 2D)**.

Having demonstrated the versatility of our platform for two cell types, we continued experiments with U2OS cells for all statistics presented in the next sections. To demonstrate that our technique can reliably control cell shape on nanopillars, we quantified the conformity of cells to micropattern shapes and compared it to micropatterns on flat substrates. Specifically, we calculated the similarity between cell shape and micropattern shape on both nanopillars and flat substrates. We used the FITC-gelatin/FN to determine contours of micropattern shapes, and F-actin for determining cell boundaries (**Figure 2E**).

Additionally, we used the Hu Moments method to determine the similarity of the ECM micropattern and cell shape as detailed in the methods section.^40^ The similarity values range from 0 to 1, where 1 indicates highest value for the similarity that demonstrates the highest conformity of cell to the micropattern shape. Our results show that there are no significant differences in the similarity values for the U2OS cells on flat and nanopillar substrates for any of the micropattern shapes.(**Figure 2F**) However, the average similarity of the cells and micropatterns and the standard deviations are slightly higher for all geometries on flat surfaces, which can be explained by the tendency of cells to grow irregularly due to the filopodia retraction membrane protrusions around the nanostructures.^41^ Interestingly, although cells conform to micropatterns of any shape on flat substrates with comparable similarity values (*i*.*e*. the similarity values are not statistically different), on nanopillars cells exhibit lower conformity to specific micropattern shapes. We observed statistically significant difference in the cell conformity to rectangular shapes with aspect ratio of 5 and 10 compared to square shape pattens and rectangular patterns with aspect ratio of 2, indicating that cells tend to resist elongated shapes on nanopillars which is consistent with previous studies of cellular morphology behavior on nanostructures^34^. This analysis indicates that the micropatterning on nanopillar platform offers high controllability for a range of geometries in single cell level. For all the future experiments we analyzed cells that conformed to the micropattern shapes with a threshold similarity value above 0.6 and excluded nonconformant cells.

### Control of cell and nuclear morphology on nanopillars

To investigate the synergic effects of micropatterns and nanopillars on cellular and nuclear morphology and deformation, we compared three important shape factors: size, roundness, and shape index. A combination of these parameters can provide a comprehensive understanding of overall cell and nuclear shape in response to the external cues.

#### Cell and nuclear size

To quantify the cell and nuclear size we measured surface area of the cell and nucleus based on actin and nucleus (DAPI) staining. Therefore, we can manipulate the shape of cells without altering their size, as our study found no significant difference in surface area among various shapes **(Figure 3A)**. Expectedly, the nuclear surface area followed a similar trend as the cell surface area **(Figure 3B)**.

**Figure 3.**
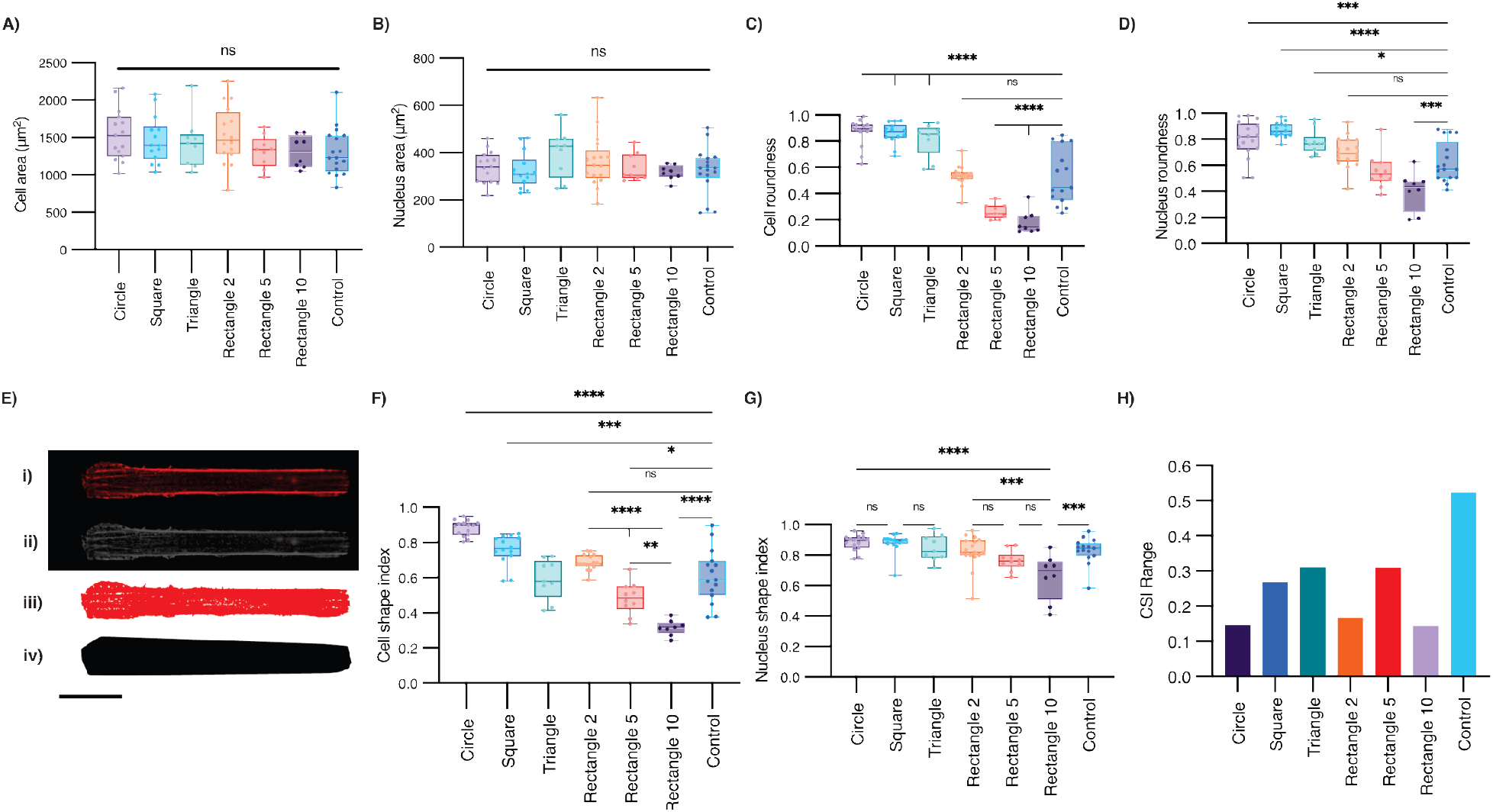
Morphology characteristic of U2OS cells and nucleus when grown on micropatterned nanopillars with various shapes and surface area of 1600 μm^2^. Comparison of the surface area of the A) U2OS cells and B) nucleus grown on micropatterns. Comparison of elongation of cells based on roundness of C) U2OS cells and D) nucleus. E) Imaging process of the cell shape analysis with ImageJ software. i) Images of cells stained with actin obtained, ii) the images converted to greyscale, iii) thresholding used to separate the cell and the background and iv) convex hull used to analyze the convex to analyze cell and nucleus shape index. Comparison of the F) Cell Shape index (CSI) of U2OS cells and G) Nucleus Cell Shape Index (NSI) on different micropatterns. H) CSI spread of the cells compared with subtracting the lowest CSI from the highest CSI within each shape. (*p < 0.05, **p < 0.01, ***p < 0.001, and ****p < 0.0001.)

#### Cell and nuclear roundness

Next, we characterized the cell roundness and elongation, as a parameter that is critical to electrical and mechanical characteristics of cells such as heart and neuronal cells.^42^ For example, Kim *et al*. studied the relationship of cell morphology on action potential and cell-cell interactions of heart muscle cells^43^. Hence, we performed roundness analysis of the cell and nucleus on micropatterns. Roundness is defined as the ratio of minor axis to the major axis of the cell. For instance, a perfect circle, triangle, and square have roundness equal to 1 while rectangle 2, rectangle 5 and rectangle 10 have roundness values equal to 0.5, 0.25, 0.1 respectively. We observed that the roundness of cells varies significantly on different geometries of ECM protein patterns. While the cells had obvious variation in roundness when spread on other shapes, rectangular shape with an aspect ratio of 2 had the highest similarity to the control cells (non-patterned) (**Figure 3C**). The nuclear deformation on micropatterns also exhibited a similar trend as the cellular response, with the rectangular shapes with aspect ratios of 2 and 5 having no significant difference to the control cells (**Figure 3D**). This can be explained by the higher mechanical modulus and stiffness of the nucleus compared to the whole cells, which makes the nuclear deformation and conformity more difficult than cellular deformation^44,45^. The combination of the cell and nuclear deformation on micropatterns reveal a native shape of U2OS cells best matching a micropattern aspect ratio of 2.

#### Cell and nuclear shape indices

To measure the cell shape index, we used the cell shape index (CSI) and nuclear shape index (NSI) (see Methods section). Specifically, for the CSI classical image analysis was not sufficient to analyze the CSI due to the membrane protrusions such as lamellipodia on epithelial cells. Hence, we developed an automated cellular analysis based on actin staining of the cells. We have optimized the smallest fitting convex hull that covers the shape to eradicate the effect of cellular irregularity **(Figure 3E)**. Our analysis showed that CSI of cells complies with the protein patterned shapes for various shapes. The average CSI values for the cells measured were 0.9, 0.76, 0.60, 0.69, 0.48, 0.31 for the circle, square, triangle, rectangle 2, rectangle 5, rectangle 10 shapes, respectively, showing minimal deviation from the protein patterned shape index (1, 0.79, 0.6, 0.7, 0.44, 0.26 for the circle, square, triangle, rectangle 2, rectangle 5, rectangle 10 shapes, respectively) as shown in **Figure 3F**. Our results also show that control non-patterned cells exhibit a wide range of cell shapes on nanopillars (CSI range of ∼0.4-0.9) (**Figure 3F-control**). Therefore, our various engineered micropattern shapes can precisely control the cell shape within the 0.4 - 0.9 control range (square, triangle, rectangle2 and rectangle5) or into new shapes that cells don’t naturally adapt if not patterned (circle or rectangle 10). This is evident from the statistically significant difference between CSI of nonpatterned control cells and cells on circle (P value of 0.001) and rectangle 10 (P value of 0.0001) micropatterns. Additionally, the difference in CSI of non-patterned control cells were statistically insignificant from the CSI of cells on square, triangle, rectangle2 and rectangle5 micropatterns.

Interestingly, despite significant changes in cell shape on various micropattern shapes, the nuclear shape was only significantly different when comparing control non-patterned cells, to highly elongated rectangle 10 patterns (NSI of 0.65 for rectangle 10 compared to NSI of 0.83 for non-patterned) suggesting that the small width of the patterns impacts nuclear shape changes (**Figure 3G**). To demonstrate the controllability of cellular shape on nanopillars using our technique, we compare the range of cell shapes on nanopillars with and without micropatterning. Our results shows that the micropatterning technique on nanopillars decreased the CSI range thoroughly across all geometries. (**Figure 3H)**

### Comparing cellular morphologies on micropatterned flat and nanopillar substrates

In the previous section, we demonstrated that cells can be confined to specific ECM shapes by comparing properties of cells on ECM-micropatterned and non-patterned nanopillars. In this section, we investigated the changes in cell and nuclear morphology when cells were confined to micropatterns on nanopillar or flat substrates. The confocal image shown in **Figure 4A** of a cell confined partially to nanotopography and partially to a flat substrate provides a clear visual representation of the complex interplay between the cells and the substrate. By controlling cell shapes on both nanopillar and flat substrates we were able to better determine how nanopillars affected cell and nuclear shape parameters, and the coupling between the cell and nuclear shape in a confined 2D ECM. Comparing the cell area on flat and nanopillar surfaces we found no notable differences, demonstrating that the controlled area of the cells on micropatterns was unaffected by the substrate shape. **(Figure 4B)** Interestingly, rectangular micropatterns with an aspect ratio of 2 manifested a difference in nuclear size between nanopillar and flat substrate, whereas the size was comparable on other micropattern shapes. (**Figure 4C**) We also compared the difference in cellular roundness between nanopillars and flat substrate. (**Figure 4D-4E**) While the effect of most shapes was independent of the nanotopography, interestingly, rectangle 2 showed a significant difference for the nuclear roundness (p < 0.0001). These findings suggest that nanopillars may alter nuclear morphology only for particular cell shapes which could be related to how the cytoskeletal elements involved in nuclear shape are organized on nanopillars for these cell shapes. Although we observed this phenomenon on rectangle 2, further studies would be required to see how different cell types and nanopillar dimensions impact these results.^3447^ Additionally, results show that cells grown on nanopillars exhibit a higher shape index, indicative of a circular shape, consistent with previous reports.^34^ However, when micropatterned ECM was used to confine the cells, no statistically significant variation in cellular and nuclear control was observed between flat and nanopillar surfaces (**Figure 4F-4G**)

**Figure 4.**
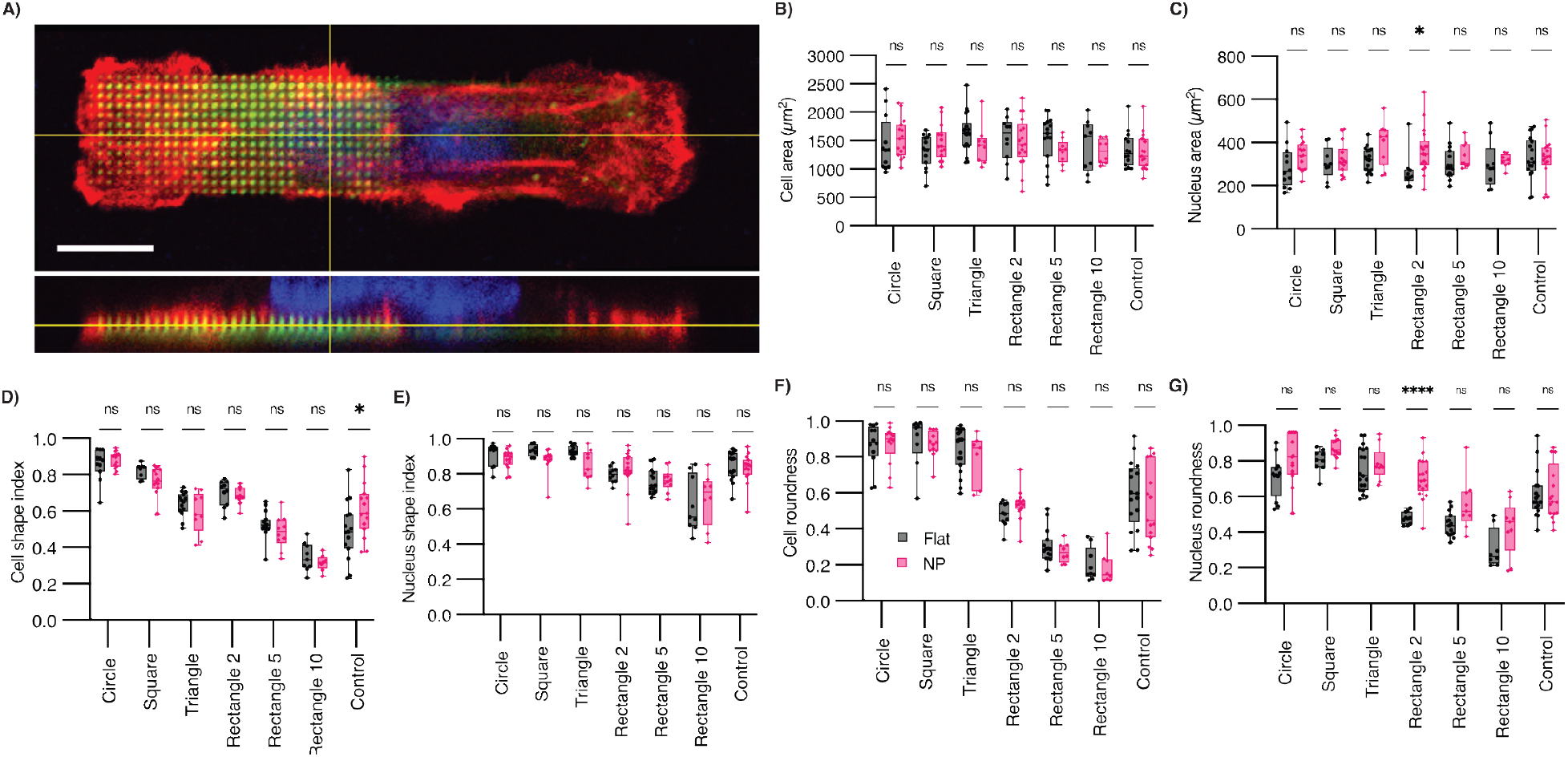
Comparisons of cell and nuclear morphology on micropatterned nanopillars and micrpatterend flat surface. A) representative confocal top and side view images of a U2OS cell growing partially on flat and nanopillar area. ECM protein gelatin (green), actin (red) and nucleus (blue). Comparison of the B) cell area and C) nucleus area for cells growing on flat and nanopillar substrate. Comparison of the D) cell and E) nucleus roundness of the U2OS cells growing on flat and nanopillar substrate. F) Cell and G) nucleus shape index comparison of U2OS cells growing on flat and nanopillar substrates. (*p < 0.05, **p < 0.01, ***p < 0.001, and ****p < 0.0001.)

Earlier studies of micropatterned cells on flat substrates have shown that the shape and orientation of the nucleus is coupled with that of the cell.^25^ Hence, some studies have used cell micropatterning as a technique to deform the nucleus.^46^ We next determined whether nanopillars affect the coupling between cell shape and nuclear shape and orientation, and whether the nucleus can be deformed by micropatterning on nanopillars. To do this, we compared the ratio of NSI to CSI on nanopillars vs flat substrates. This ratio can be studied as an index to compare the relative deformation of the cell and nucleus, where higher NSI to CSI ratio is indicative of more deformed cell, but less deformed nucleus.^25,47^ Our results in **Figure 5A** show that there is a significant difference in NSI to CSI between cells cultured on flat substrates and nanopillars without micropatterning; however, when confining the cells with protein micropatterns there is no significant difference between flat and nanopillar substrate suggesting that the nucleus can be deformed by controlling cell shape on nanopillar substrates. However, our results show that the NSI to CSI ratio is highest for rectangle 10 suggesting that the nucleus is rounder and not deformed as much as the cell for this cell shape (**Figure S2**). ^50–52^These results suggest that the nucleus can be controllably deformed by micropatterning and engineering cell morphology on nanopillars; However, the degree of nuclear deformation is limited for high aspect ratio cells.

**Figure 5.**
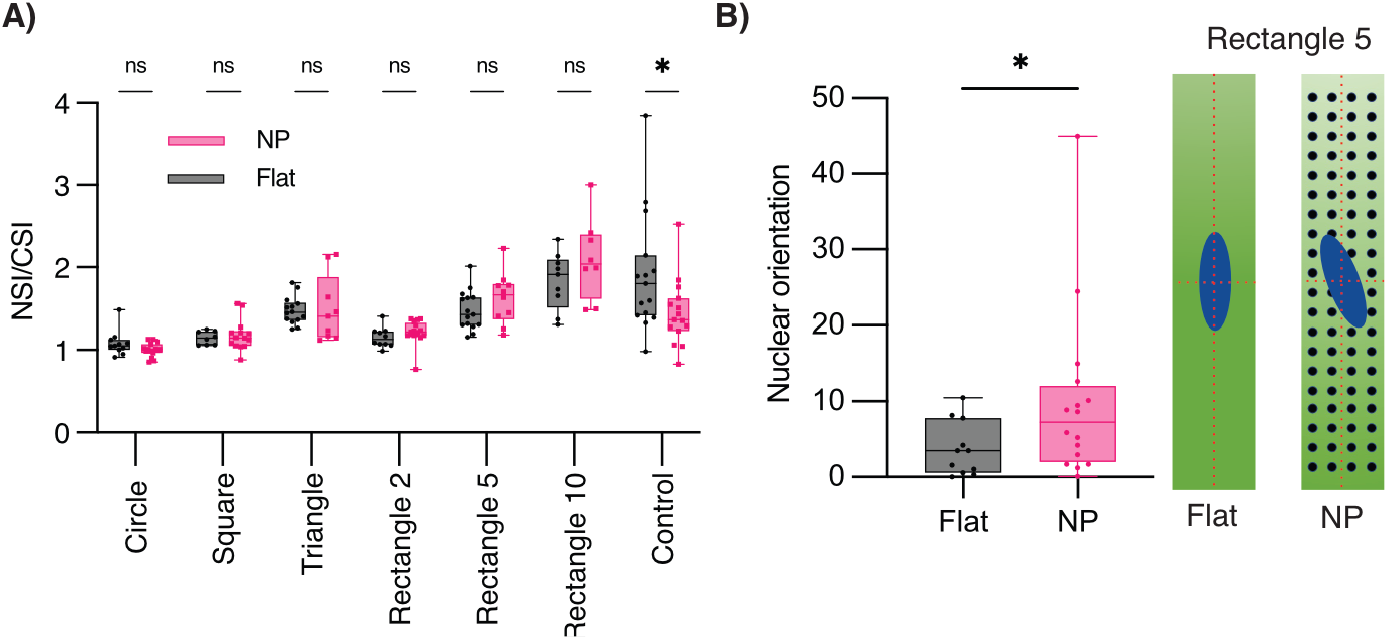
Comparison of the cell and nuclear shape coupling, and nuclear orientation on nanopillar and flat substrates. A) Ratio of nuclear shape index (NSI) to the cellular shape index (CSI) for U2OS cells on flat and nanopillar substrate, respectively. B) Changes in nuclear orientation on rectangle 5 micropatterns on nanopillars compared with corresponding micropatterns on flat substrates. (*p < 0.05)

We next asked whether nanopillars can affect nuclear orientation on micropatterns when cells are elongated. Several studies have shown that cells and nucleus reorient themselves along the direction of maximum stress to minimize the applied force. For example, cells on the elongated shapes align themselves along the major axis of the rectangle, which can cause the nucleus to be also aligned along the same axis.^25^ We computed and compared the nuclear orientation on rectangle 5 between flat and nanopillar substrates. The flat substrate shows absolute rotation of 3.7±3.5° while nanopillar shows 9.9±11.2° values (**Figure 5B**). These findings suggests that nanopillars can disrupt the orientation and alignment of the nucleus that is caused by cell shape. These findings and further studies using our technique can provide fundamental understanding of how nanopillars impact cell migration^48^, signaling^49^ and division^50^.

### Effect of cell shape on nanopillar-enhanced endocytosis

Nanopillars are known to enhance cell endocytosis.^28^ Previous studies have shown that especially nanoscale membrane curvature can enhance the clathrin-mediated endocytosis.^29,51^ In order to further investigate the effect of nanopillars in combination with cellular elongation, we conducted experiments using U2OS cells and FM 1-43 dye^52^ based on previously described method.^52^ FM 1-43 dye has minimal fluorescent signal when present in the solution; however, when it binds to the membrane it will be highly fluorescent.^53^ **Figure 6A** shows a schematic of the curvature-related enhancement of clathrin-mediated endocytosis and our results that nanopillars enhance the endocytosis compared to flat surface. Next, we studied the effect of elongation and cellular shape on nanopillar-enhanced endocytosis. Quantitative analysis of the endocytosis on nanopillars shows clearly that shapes with higher elongation (rectangular shapes) have more FM 1-43 dye endocytosis than circular shapes as shown in **Figure 6B**. Endocytosis is known to be regulated through the cytoskeleton, which has also been identified as the link between cell and nuclear shape regulation.^25^ Since we observed that cells on rectangle 10 exhibited the least deformed nucleus (Figure 5B), this could indicate that the increased endocytosis is related to a disrupted cytoskeletal organization. Nanotopography can likewise affect the cell cytoskeleton and endocytosis, which we believe highlights the importance of our platform for disentangling the effect of different topographical and biochemical cues.^54^ This is extremely important due to the application of nanopillars for intracellular drug delivery and sensing platforms.^55^ **Figure 6C** shows overlay images of U2OS cells on rectangular and circular shapes with the vesicles endocytosed to the cell. The results of these experiments provide insights into the complex nature of the cellular response to substrate and cellular elongation.

**Figure 6.**
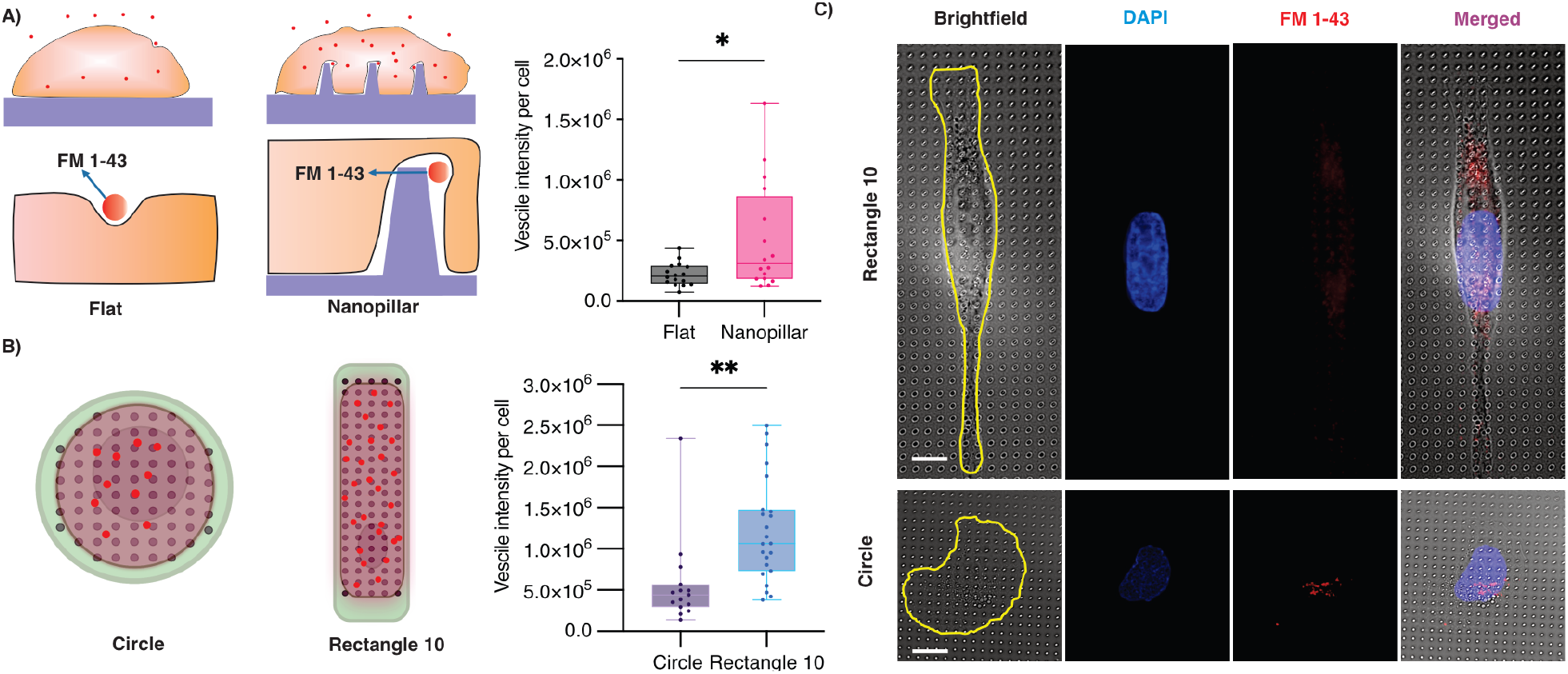
Effect of cell-shape on nanopillar-enhanced endocytosis. A) Schematic showing the proposed mechanism of endocytosis based on substrate (flat and nanopillar) and quantitative analysis of endocytosed FM 1-43 dye in U2OS cells. B) Schematic of U2OS cells growing in different geometries (circle and rectangle 10) on nanopillars and quantitative analysis of endocytosed FM 1-43 dye. C) Representative fluorescence images of the FM 1-43 dye endocytosed in U2OS cells. Brightfield image is used to analyze cell shape, FM 1-43 dye (red) and nucleus (blue). (*p < 0.05, and **p < 0.01.)

## CONCLUSION

Nanopillars are emerging as building blocks of platforms for a variety of exciting biomedical applications including fundamental electro- and mechano-biology^21^, cancer diagnostics^22^, and drug delivery and discovery^5,6,15^. Our results show that cells growing on nanopillars exhibit higher shape indices (**Figure 4F**) indicating the tendency of the cells to grow to the more circular shape on our nanopillars than flat substrates, which agrees with previous results by Modaresifar *et al*.^34^ In this work we show that maskless and contact-less protein micropatterning can be used to modulate and regulate cellular shape on flat and nanopillar substrates. By direct comparison of cells with similar shapes on nanopillar vs flat substrates, we show that the nucleus can be deformed by controlling cell shape on both substrates. However, our results show that nuclear orientation along the major cell axis is disrupted on nanopillars. Moreover, we show that nanopillar-enhanced endocytosis can be further increased if cells are elongated on nanopillars (**Figure 6B**). Since cell shape can also be controlled by nanopillar geometry and spacing^56^, these findings have significant implications for the development of nanostructured platforms for biomolecule delivery.

## EXPERIMENTAL METHODS

### Nanopillar fabrication

Nanopillar fabrication started with procurement of 4-inch fused quartz wafer from Wafer Pro (USA). The wafer underwent RCA cleaning using SC1 and SC3, followed by spin rinsing and drying in a SRD (MEI). Subsequently, they were then spin-coated with AZ 1512 photoresist (EMD Performance Materials Corp., USA) at 4000 rpm for 45 seconds at 12000 acc using a 3-step process, resulting in a thickness of approximately 1.2 um. The wafers were patterned and exposed (375nm, dose 300) using our design file (.gds) in the Heidelberg MLA system. After exposure, the substrate were developed with AZ400 (AZ Electronic Materials USA Corp) for 30 seconds and checked under an optical microscope. The Cr (99.998% target) metal was deposited onto the patterned substrate using a Temescal Ebeam evaporator, followed by a lift-off procedure using RR41, acetone, and IPA respectively. The substrates were then dry etched for 50 minutes using the Oxford Plasmalab 80plus RIE system with Ar (35 sccm) and ChF3 (25 sccm) gases at 50mT and 200W to etch the wafer and create nanopillars. Finally, the substrates underwent wet etching for 10 minutes using Cr etchant (Transene Company Inc.) to remove the Cr, followed by BOE 20:1 for 16 minutes to remove the exposed quarts. The wafers were diced into 1 cm x 1 cm chips for further experiments.

### Micropatterning method

Maskless lithography method with PRIMO were done based on described method^37^ with modifications for patterning on nanopillar surface. Briefly, nanopillars surface from quartz were UVO treated for 10 minutes to clean the surface and enhance the electrostatic adsorption of positively charge PLL (Sigma Aldrich, USA) groups to the negatively charged surface. PLL with 0.1 mg/L were incubated on the surface for 30 minutes and washed with 0.1 M HEPES (Gibco ™, USA) buffer. Following PLL incubation, m-PEG-SVA (Laysan Bio, USA) with the concentration of 100 mg/mL were prepared and added to the surface and incubated for 1 hr. The resulting PEG layer serves as an antifouling layer that can hinder protein and cellular attachment. The chips were washed for 10 times with DI water and air-dried. The UV sensitive PLPP gel (Alvéole, Fr) solution were mixed with 70% v/v ethanol and added to the dried surface and covered till micropatterning. The chips were mounted on a glass substrate on a stage of a Nikon Ti2 Eclipse microscope. The images were made by open-source software Inkscape and loaded into the Leonardo (Alvéole, Fr) software. The patterns were projected using a 375 nm laser on a S Plan Fluor ELWD 20x with 0.45 NA objective with various doses. The projected area causes localized cleavage of the PEG-SVA, exposing PLL to ECM protein attachment. Chips were washed with DI water 10 times and incubated with 20 μg/mL mixture of FITC-gelatin (BioVision, USA) and FN (Sigma Aldrich, USA) with 1:1 w/w? ratio. Chips were washed with 1x PBS (Gibco™, USA) 10 times profusely to remove the extra protein and prepares the sample for cell seeding.

### SEM imaging

Topography of the nanopillar arrays were characterized with scanning electron microscopy (Quanta FEG 250, FEI), under low vacuum. Images were obtained at 5 KV with Everhart Thornley detector (ETD). Images were taken under 45 degrees tilted for height visualization.

### Cell culture and seeding

U2OS(ATCC) cells were grown in McCoy’s 5A Medium (ATCC) supplemented with 10% (vol/vol) fetal bovine serum (FBS)(Sigma Aldrich, USA) and 1% (vol/vol) penicillin-streptomycin (Sigma-Aldrich, USA) and incubated at 37 °C with 5% CO2. U2OS cells at passage 2-8 were detached by TrypLE™ Express Enzyme (1X) (Gibco™, USA). After centrifugation of the cell suspension, supernatant removed from to achieve the cell pellet. Cell pellet were resuspended in the McCoy’s 5a media and the abovementioned supplements. A 100 μL cell suspension with ∼ 1×10^4^ was added to each nanopillar chips and incubated at 37 °C. Cells were washed after 30 mins after initial adhesion to the ECM micropatterns to achieve single cells per patterns and media added the chip and incubated for growth. Cells were fixed after 24 h and used for further analysis.

### Fluorescent staining and microscopy

After 24 h, cells were fixed with 4% paraformaldehyde (Electron Microscopy Sciences, USA) at room temperature for 10 mins and washed with PBS. Cells were permeabilized with 1% with Triton^TM^ X-100 (Sigma-Aldrich, USA) for 10 minutes and blocked in 2% (wt/vol) Bovine Serum Albumin (BSA)(Thermo Scientific™, USA) for 1 h. Sample were rinsed in PBS and incubated with 4’,6-diamidino-2-phenylindole (DAPI) (Thermo Scientific™, USA) for 5 mins and Alexa 594-phalloidin (Invitrogen™, USA) for 20 mins in dark for the nucleus and F-actin staining respectively. Images were collected with Echo revolve microscope with 60x PLAN Fluorite water dipping objective with NA 1.00.

### Method for the cell analysis

The image analysis for the fluorescent images and morphology analysis performed using ImageJ 1.53 (NIH, US). The signal intensity of the fluorescent labeled micropatterns measured by selecting the shapes in grayscale and calculating grey value of shapes after applying subtract background by choosing sliding paraboloid and enhancing contrast of the images by CLAHE by block size of 19, histogram bins of 256 with maximum slope of 3. Additionally, the point-by-point signal of micropatterning measured by using plot profile command as described elsewhere.^57^ The area and roundness of cell and nucleus was calculated by analyze particle command as follows:

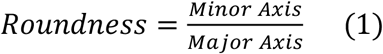

The Cellular Shape Index (CSI) calculate based on actin staining to achieve the area and perimeters of the cell as follows.

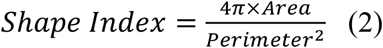

To remove the intensity dependency of analysis for the shape analysis we have used the convexHull command of OpenCV library using Python by drawing the smallest counters surrounding the cells to achieve more accurate values for cell and nucleus shape index (CSI and NSI) and rely solely on the shape of the objects.

To investigate the conformity of the cells to the micropatterned proteins as an indicator of micropatterning effectiveness we have implemented Hu Moments for shape analysis of two objects. Hu Moments are a set of mathematical descriptors that can be used for shape analysis of objects while they are invariant to image transformations that allows Hu Moments to provide a reliable and consistent measurement of shape analysis. Raw image moment can be calculated using equation (1) while the I (x,y) is the intensity of the pixel at given (x, y) location.^58^

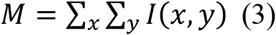

### Endocytosis

For the endocytosis analysis, U2OS cells were cultures on various substrates after 24 h of incubation. media removed following by washing the cells with PBS. FM 1-43 (ThermoFisher, USA) with 2μM concentration added to the well-plate containing nanopillar chip. Cells were incubated for 15 minutes with media containing FM 1-43 dye in 37°C. Cells were washed with PBS and incubated for additional 20 mins in 37 °C. Cells were fixed with 4% PFA for 10 mins and stained with 300 nM DAPI for 5 mins. The sample were imaged and analyzed using ImageJ (NIH, USA) software.

### Statistical analysis

All the data shown as the mean ± standard deviation (SD). Statistical significance with two-tailed Student’s t-test analysis with Welch’s correction was performed in Prism (GraphPad, USA) to determine the statistical significance of the differences between the means of the different experimental groups. In addition, the results of figure 4 for comparison of the cells on flat and nanopillar within different micropattern shapes were analyzed using two-way ANOVA, followed by Tukey’s multiple comparisons post hoc analysis.

## ACKNOWLEDGMENT

This work was performed in part at the San Diego Nanotechnology Infrastructure (SDNI) of UCSD, a member of the National Nanotechnology Coordinated Infrastructure, which is supported by the National Science Foundation (Grant ECCS-2025752). This work was in part supported by Air Force Office of Scientific Research YIP award (Award Number: 311616-00001) and Cancer research coordinating committee faculty seed grant to ZJ. LHK acknowledges support from the the Carlsberg Foundation (CF19-0742).

## REFERENCES

(1) Milos, F.; Belu, A.; Mayer, D.; Maybeck, V.; Offenhäusser, A. Polymer Nanopillars Induce Increased Paxillin Adhesion Assembly and Promote Axon Growth in Primary Cortical Neurons. Adv Biol 2021, 5 (2), 2000248.

(2) Sorgato, M.; Guidi, E.; Conconi, M. T.; Lucchetta, G. Surface Nanostructuring of Bioresorbable Implants to Induce Osteogenic Differentiation of Human Mesenchymal Stromal Cells. CIRP Annals 2021, 70 (1), 463–466.

(3) Ghosh, L. das; Hasan, J.; Jain, A.; Sundaresan, N. R.; Chatterjee, K. A Nanopillar Array on Black Titanium Prepared by Reactive Ion Etching Augments Cardiomyogenic Commitment of Stem Cells. Nanoscale 2019, 11 (43), 20766–20776.

(4) He, G.; Chen, H.; Liu, D.; Feng, Y.; Yang, C.; Hang, T.; Wu, J.; Cao, Y.; Xie, X. Fabrication of Various Structures of Nanostraw Arrays and Their Applications in Gene Delivery. Adv Mater Interfaces 2018, 5 (10), 1701535.

(5) Stewart, M. P.; Sharei, A.; Ding, X.; Sahay, G.; Langer, R.; Jensen, K. F. In Vitro and Ex Vivo Strategies for Intracellular Delivery. Nature 2016, 538 (7624), 183–192.

(6) Ma, Y.; Nolte, R. J. M.; Cornelissen, J. J. L. M. Virus-Based Nanocarriers for Drug Delivery. Adv Drug Deliv Rev 2012, 64 (9), 811–825.

(7) Lee, W. S.; Ahn, J.; Jung, S.; Lee, J.; Kang, T.; Jeong, J. Biomimetic Nanopillar-Based Biosensor for Label-Free Detection of Influenza A Virus. Biochip J 2021, 15 (3), 260–267.

(8) Cui, S.; Tian, C.; Mao, J.; Wu, W.; Fu, Y. Nanopillar Array-Based Plasmonic Metasurface for Switchable Multifunctional Biosensing. Opt Commun 2022, 506, 127548.

(9) Losero, E.; Jagannath, S.; Pezzoli, M.; Lashuel, H. A.; Galland, C.; Quack, N. Neuronal Growth on High-Aspect-Ratio Diamond Nanopillar Arrays for Biosensing Applications. arXiv preprint arXiv:2207.09903 /i>2022.

(10) Jahed, Z.; Yang, Y.; Tsai, C.-T.; Foster, E. P.; McGuire, A. F.; Yang, H.; Liu, A.; Forro, C.; Yan, Z.; Jiang, X. Nanocrown Electrodes for Parallel and Robust Intracellular Recording of Cardiomyocytes. Nat Commun 2022, 13 (1), 2253.

(11) Li, X.; Matino, L.; Zhang, W.; Klausen, L.; McGuire, A. F.; Lubrano, C.; Zhao, W.; Santoro, F.; Cui, B. A Nanostructure Platform for Live-Cell Manipulation of Membrane Curvature. Nat Protoc 2019, 14 (6), 1772–1802.

(12) Elnathan, R.; Tay, A.; Voelcker, N. H.; Chiappini, C. The Start-Ups Taking Nanoneedles into the Clinic. Nat Nanotechnol 2022. https://doi.org/10.1038/s41565-022-01158-5.

(13) Hanson, L.; Zhao, W.; Lou, H.-Y.; Lin, Z. C.; Lee, S. W.; Chowdary, P.; Cui, Y.; Cui, B. Vertical Nanopillars for in Situ Probing of Nuclear Mechanics in Adherent Cells. Nat Nanotechnol 2015, 10 (6), 554–562.

(14) Finbloom, J. A.; Huynh, C.; Huang, X.; Desai, T. A. Bioinspired Nanotopographical Design of Drug Delivery Systems. Nature Reviews Bioengineering 2023, 1 (2), 139–152. https://doi.org/10.1038/s44222-022-00010-8.

(15) Xie, X.; Xu, A. M.; Leal-Ortiz, S.; Cao, Y.; Garner, C. C.; Melosh, N. A. Nanostraw– Electroporation System for Highly Efficient Intracellular Delivery and Transfection. ACS Nano 2013, 7 (5), 4351–4358. https://doi.org/10.1021/nn400874a.

(16) Shokoohimehr, P.; Cepkenovic, B.; Milos, F.; Bednár, J.; Hassani, H.; Maybeck, V.; Offenhäusser, A. High-Aspect-Ratio Nanoelectrodes Enable Long-Term Recordings of Neuronal Signals with Subthreshold Resolution. Small 2022, 18 (22), 2200053. https://doi.org/10.1002/smll.202200053.

(17) Losero, E.; Jagannath, S.; Pezzoli, M.; Goblot, V.; Babashah, H.; Lashuel, H. A.; Galland, C.; Quack, N. Neuronal Growth on High-Aspect-Ratio Diamond Nanopillar Arrays for Biosensing Applications. Sci Rep 2023, 13 (1), 5909. https://doi.org/10.1038/s41598-023-32235-x.

(18) Xie, C.; Lin, Z.; Hanson, L.; Cui, Y.; Cui, B. Intracellular Recording of Action Potentials by Nanopillar Electroporation. Nat Nanotechnol 2012, 7 (3), 185–190. https://doi.org/10.1038/nnano.2012.8.

(19) Fang, J.; Xu, D.; Wang, H.; Wu, J.; Li, Y.; Yang, T.; Liu, C.; Hu, N. Scalable and Robust Hollow Nanopillar Electrode for Enhanced Intracellular Action Potential Recording. Nano Lett 2023, 23 (1), 243–251. https://doi.org/10.1021/acs.nanolett.2c04222.

(20) Lin, Z. C.; Xie, C.; Osakada, Y.; Cui, Y.; Cui, B. Iridium Oxide Nanotube Electrodes for Sensitive and Prolonged Intracellular Measurement of Action Potentials. Nat Commun 2014, 5 (1), 3206. https://doi.org/10.1038/ncomms4206.

(21) Jahed, Z.; Yang, Y.; Tsai, C.-T.; Foster, E. P.; McGuire, A. F.; Yang, H.; Liu, A.; Forro, C.; Yan, Z.; Jiang, X.; Zhao, M.-T.; Zhang, W.; Li, X.; Li, T.; Pawlosky, A.; Wu, J. C.; Cui, B. Nanocrown Electrodes for Parallel and Robust Intracellular Recording of Cardiomyocytes. Nat Commun 2022, 13 (1), 2253. https://doi.org/10.1038/s41467-022-29726-2.

(22) Zeng, Y.; Zhuang, Y.; Vinod, B.; Guo, X.; Mitra, A.; Chen, P.; Saggio, I.; Shivashankar, G. V.; Gao, W.; Zhao, W. Guiding Irregular Nuclear Morphology on Nanopillar Arrays for Malignancy Differentiation in Tumor Cells. Nano Lett 2022, 22 (18), 7724–7733. https://doi.org/10.1021/acs.nanolett.2c01849.

(23) Li, N.; Jin, K.; Chen, T.; Li, X. A Static Force Model to Analyze the Nuclear Deformation on Cell Adhesion to Vertical Nanostructures. Soft Matter 2022, 18 (35), 6638–6644. https://doi.org/10.1039/D2SM00971D.

(24) Capozza, R.; Caprettini, V.; Gonano, C. A.; Bosca, A.; Moia, F.; Santoro, F.; De Angelis, F. Cell Membrane Disruption by Vertical Micro-/Nanopillars: Role of Membrane Bending and Traction Forces. ACS Appl Mater Interfaces 2018, 10 (34), 29107–29114. https://doi.org/10.1021/acsami.8b08218.

(25) Versaevel, M.; Grevesse, T.; Gabriele, S. Spatial Coordination between Cell and Nuclear Shape within Micropatterned Endothelial Cells. Nat Commun 2012, 3 (1), 671. https://doi.org/10.1038/ncomms1668.

(26) Hanson, L.; Lin, Z. C.; Xie, C.; Cui, Y.; Cui, B. Characterization of the Cell–Nanopillar Interface by Transmission Electron Microscopy. Nano Lett 2012, 12 (11), 5815–5820. https://doi.org/10.1021/nl303163y.

(27) Nakamoto, M. L.; Forró, C.; Zhang, W.; Tsai, C.-T.; Cui, B. Expansion Microscopy for Imaging the Cell–Material Interface. ACS Nano 2022, 16 (5), 7559–7571. https://doi.org/10.1021/acsnano.1c11015.

(28) Li, X.; Klausen, L. H.; Zhang, W.; Jahed, Z.; Tsai, C.-T.; Li, T. L.; Cui, B. Nanoscale Surface Topography Reduces Focal Adhesions and Cell Stiffness by Enhancing Integrin Endocytosis. Nano Lett 2021, 21 (19), 8518–8526. https://doi.org/10.1021/acs.nanolett.1c01934.

(29) Zhao, W.; Hanson, L.; Lou, H.-Y.; Akamatsu, M.; Chowdary, P. D.; Santoro, F.; Marks, J. R.; Grassart, A.; Drubin, D. G.; Cui, Y.; Cui, B. Nanoscale Manipulation of Membrane Curvature for Probing Endocytosis in Live Cells. Nat Nanotechnol 2017, 12 (8), 750–756. https://doi.org/10.1038/nnano.2017.98.

(30) Teo, B. K. K.; Goh, S.-H.; Kustandi, T. S.; Loh, W. W.; Low, H. Y.; Yim, E. K. F. The Effect of Micro and Nanotopography on Endocytosis in Drug and Gene Delivery Systems. Biomaterials 2011, 32 (36), 9866–9875. https://doi.org/10.1016/j.biomaterials.2011.08.088.

(31) Zhou, J.; Zhang, X.; Sun, J.; Dang, Z.; Li, J.; Li, X.; Chen, T. The Effects of Surface Topography of Nanostructure Arrays on Cell Adhesion. Physical Chemistry Chemical Physics 2018, 20 (35), 22946–22951. https://doi.org/10.1039/C8CP03538E.

(32) Beckwith, K. S.; Ullmann, S.; Vinje, J.; Sikorski, P. Influence of Nanopillar Arrays on Fibroblast Motility, Adhesion, and Migration Mechanisms. Small 2019, 15 (43), 1902514. https://doi.org/10.1002/smll.201902514.

(33) Wang, K.; Man, K.; Liu, J.; Meckes, B.; Yang, Y. Dissecting Physical and Biochemical Effects in Nanotopographical Regulation of Cell Behavior. ACS Nano 2023, 17 (3), 2124–2133. https://doi.org/10.1021/acsnano.2c08075.

(34) Modaresifar, K.; Ganjian, M.; Díaz-Payno, P. J.; Klimopoulou, M.; Koedam, M.; van der Eerden, B. C. J.; Fratila-Apachitei, L. E.; Zadpoor, A. A. Mechanotransduction in High Aspect Ratio Nanostructured Meta-Biomaterials: The Role of Cell Adhesion, Contractility, and Transcriptional Factors. Mater Today Bio 2022, 16, 100448.

(35) Lestrell, E.; Chen, Y.; Aslanoglou, S.; O’Brien, C. M.; Elnathan, R.; Voelcker, N. H. Silicon Nanoneedle-Induced Nuclear Deformation: Implications for Human Somatic and Stem Cell Nuclear Mechanics. ACS Appl Mater Interfaces 2022, 14 (40), 45124–45136. https://doi.org/10.1021/acsami.2c10583.

(36) Yang, L.; Jurczak, K. M.; Ge, L.; Rijn, P. High-Throughput Screening and Hierarchical Topography-Mediated Neural Differentiation of Mesenchymal Stem Cells. Adv Healthc Mater 2020, 9 (11), 2000117. https://doi.org/10.1002/adhm.202000117.

(37) Strale, P.-O.; Azioune, A.; Bugnicourt, G.; Lecomte, Y.; Chahid, M.; Studer, V. Multiprotein Printing by Light-Induced Molecular Adsorption. Advanced Materials 2016, 28 (10), 2024–2029. https://doi.org/10.1002/adma.201504154.

(38) Melero, C.; Kolmogorova, A.; Atherton, P.; Derby, B.; Reid, A.; Jansen, K.; Ballestrem, C. Light-Induced Molecular Adsorption of Proteins Using the PRIMO System for Micro-Patterning to Study Cell Responses to Extracellular Matrix Proteins. JoVE (Journal of Visualized Experiments) 2019, No. 152, e60092.

(39) Toro-Nahuelpan, M.; Zagoriy, I.; Senger, F.; Blanchoin, L.; Théry, M.; Mahamid, J. Tailoring Cryo-Electron Microscopy Grids by Photo-Micropatterning for in-Cell Structural Studies. Nat Methods 2020, 17 (1), 50–54.

(40) žunić, D.; žunić, J. Shape Ellipticity Based on the First Hu Moment Invariant. Inf Process Lett 2013, 113 (19–21), 807–810. https://doi.org/10.1016/j.ipl.2013.07.020.

(41) Jahed, Z.; Molladavoodi, S.; Seo, B. B.; Gorbet, M.; Tsui, T. Y.; Mofrad, M. R. K. Cell Responses to Metallic Nanostructure Arrays with Complex Geometries. Biomaterials 2014, 35 (34), 9363–9371. https://doi.org/10.1016/j.biomaterials.2014.07.022.

(42) Aubin, H.; Nichol, J. W.; Hutson, C. B.; Bae, H.; Sieminski, A. L.; Cropek, D. M.; Akhyari, P.; Khademhosseini, A. Directed 3D Cell Alignment and Elongation in Microengineered Hydrogels. Biomaterials 2010, 31 (27), 6941–6951.

(43) Kim, D.-H.; Lipke, E. A.; Kim, P.; Cheong, R.; Thompson, S.; Delannoy, M.; Suh, K.-Y.; Tung, L.; Levchenko, A. Nanoscale Cues Regulate the Structure and Function of Macroscopic Cardiac Tissue Constructs. Proceedings of the National Academy of Sciences 2010, 107 (2), 565–570.

(44) Zhang, J.; Alisafaei, F.; Nikolić, M.; Nou, X. A.; Kim, H.; Shenoy, V. B.; Scarcelli, G. Nuclear Mechanics within Intact Cells Is Regulated by Cytoskeletal Network and Internal Nanostructures. Small 2020, 16 (18), 1907688. https://doi.org/10.1002/smll.201907688.

(45) Barreto, S.; Clausen, C. H.; Perrault, C. M.; Fletcher, D. A.; Lacroix, D. A Multi-Structural Single Cell Model of Force-Induced Interactions of Cytoskeletal Components. Biomaterials 2013, 34 (26), 6119–6126. https://doi.org/10.1016/j.biomaterials.2013.04.022.

(46) Bautista, M.; Fernandez, A.; Pinaud, F. A Micropatterning Strategy to Study Nuclear Mechanotransduction in Cells. Micromachines (Basel) 2019, 10 (12), 810. https://doi.org/10.3390/mi10120810.

(47) Liu, R.; Liu, Q.; Pan, Z.; Liu, X.; Ding, J. Cell Type and Nuclear Size Dependence of the Nuclear Deformation of Cells on a Micropillar Array. Langmuir 2019, 35 (23), 7469–7477. https://doi.org/10.1021/acs.langmuir.8b02510.

(48) Maninová, M.; Iwanicki, M. P.; Vomastek, T. Emerging Role for Nuclear Rotation and Orientation in Cell Migration. Cell Adh Migr 2014, 8 (1), 42–48. https://doi.org/10.4161/cam.27761.

(49) Gundersen, G. G.; Worman, H. J. Nuclear Positioning. Cell 2013, 152 (6), 1376–1389. https://doi.org/10.1016/j.cell.2013.02.031.

(50) Gudipaty, S. A.; Lindblom, J.; Loftus, P. D.; Redd, M. J.; Edes, K.; Davey, C. F.; Krishnegowda, V.; Rosenblatt, J. Mechanical Stretch Triggers Rapid Epithelial Cell Division through Piezo1. Nature 2017, 543 (7643), 118–121. https://doi.org/10.1038/nature21407.

(51) Akamatsu, M.; Vasan, R.; Serwas, D.; Ferrin, M. A.; Rangamani, P.; Drubin, D. G. Principles of Self-Organization and Load Adaptation by the Actin Cytoskeleton during Clathrin-Mediated Endocytosis. Elife 2020, 9. https://doi.org/10.7554/eLife.49840.

(52) Li, X.; Klausen, L. H.; Zhang, W.; Jahed, Z.; Tsai, C.-T.; Li, T. L.; Cui, B. Nanoscale Surface Topography Reduces Focal Adhesions and Cell Stiffness by Enhancing Integrin Endocytosis. Nano Lett 2021, 21 (19), 8518–8526.

(53) Bertrand, C. A.; Laboisse, C.; Hopfer, U.; Bridges, R. J.; Frizzell, R. A. Methods for Detecting Internalized, FM 1-43 Stained Particles in Epithelial Cells and Monolayers. Biophys J 2006, 91 (10), 3872–3883. https://doi.org/10.1529/biophysj.106.086983.

(54) Lou, H.-Y.; Zhao, W.; Zeng, Y.; Cui, B. The Role of Membrane Curvature in Nanoscale Topography-Induced Intracellular Signaling. Acc Chem Res 2018, 51 (5), 1046–1053. https://doi.org/10.1021/acs.accounts.7b00594.

(55) Chen, N.; He, Y.; Zang, M.; Zhang, Y.; Lu, H.; Zhao, Q.; Wang, S.; Gao, Y. Approaches and Materials for Endocytosis-Independent Intracellular Delivery of Proteins. Biomaterials 2022, 286, 121567. https://doi.org/10.1016/j.biomaterials.2022.121567.

(56) Vinje, J. B.; Guadagno, N. A.; Progida, C.; Sikorski, P. Analysis of Actin and Focal Adhesion Organisation in U2OS Cells on Polymer Nanostructures. Nanoscale Res Lett 2021, 16 (1), 143. https://doi.org/10.1186/s11671-021-03598-9.

(57) Horzum, U.; Ozdil, B.; Pesen-Okvur, D. Step-by-Step Quantitative Analysis of Focal Adhesions. MethodsX 2014, 1, 56–59. https://doi.org/10.1016/j.mex.2014.06.004.

(58) Körtgen, M.; Park, G.-J.; Novotni, M.; Klein, R. 3D Shape Matching with 3D Shape Contexts. In The 7th central European seminar on computer graphics; Budmerice Slovakia, 2003; Vol. 3, pp 5–17.

